# The genetical evolution of social preferences: where the categorical imperatives of Hamilton, Kant and Nash meet

**DOI:** 10.1101/2025.09.24.678278

**Authors:** Laurent Lehmann

## Abstract

This paper models the genetical evolution of individual behavioral rules that guide the choice of strategies in pairwise assortative interactions under incomplete information. Building on results at the crossroads of evolutionary theory and game theory, it is first shown that in an uninvadable population state of behavioral rule evolution, individuals are compelled to use strategies that are Nash equilibria of a lineage fitness game. Thus, choice behavior evolves to be representable as the maximization of a utility function, as if each individual holds a personal preference that orders both their own and their interaction partner’s strategies. Second, the paper contrasts two representations of personal utility that are found to be uninvadable. The first is semi-Kantian in form. This preference averages a fitness self-interest with a relatedness weighted Kantian interest. The latter interest evaluates the consequence of own behavior for own fitness, assuming the interaction partner adopts the same behavior as self. The second preference is a personal inclusive fitness. This preference combines a self-regarding interest with a relatedness weighted other-regarding interest. Each such interest takes the form of an average effect, which evaluates the consequence of expressing own behavior, instead of average population behavior, on a statistical average fitness to self and the interaction partner.

Species following the model should tend to evolve behavior such that each organism appears to be attempting to maximize its inclusive fitness.

W.D. Hamilton (1964)

## 1 Introduction

The preceding quote, taken from the abstract of Hamilton (1964), has been instrumental in promoting the view that natural selection shapes an organism’s phenotype to maximize a specific objective: inclusive fitness. This view can be understood to embody two key features about the evolutionary process. First, adaptation results from differences in fitness. However, this fitness is not that of the group, nor even that of the individual, but that of the gene lineage, since the Mendelized evolutionary process is fundamentally gene-centric (Hamilton, 1963; Dawkins, 1978; Akçay and Van Cleve, 2016). Second, to connect gene-centrism to the organism, one can conceive of an evolved individual as a nepotistic actor. Such an actor not only affects its own survival and reproduction, but also that of other individuals, with the evolutionary impact of actions depending, in some way or another, on the genetic relatedness between the actor and those affected (Hamilton, 1971; Frank, 1997; Rousset, 2015). This actor-centered perspective on evolved phenotypes enables one to ask some of the most provocative questions about adaptation, such as when would an individual be favored to sacrifice its own survival and reproduction to inflict even greater harm on another? Consequently, inclusive fitness maximization has become a foundational concept in behavioral ecology (Davies et al., 2012, chapter 11), where it serves as a powerful mechanism to generate predictions about the behavior of organisms, including humans and their culture (Alexander, 1979, see pp. 156–161 for a remarkable list of predictions).

Hamilton (1964), however, did not explicitly formalize phenotypes, nor did he demonstrate that natural selection drives interactive behavior to maximize any particular objective. His model addresses allele frequency change under natural selection when allelic effects on the survival and reproduction of actors and recipients are marginal (weak selection) and gene action is additive. Recast in terms of phenotypic evolution under social interactions, such changes can readily produce fitness minima under gradual evolution (e.g., Ajar, 2003; Avila and Mullon, 2023). Although incredibly wide-ranging and fundamental to evolutionary biology, it has become clear that Hamilton’s (1964) results provide only necessary conditions for evolutionary stability (e.g., Taylor, 1989; Rousset and Billiard, 2000; Rousset, 2004; Wakano et al., 2012; Van Cleve, 2015; Avila and Mullon, 2023). This leaves open the question of what behavioral objective organisms evolve to maximize, if any at all.

This paper combines arguments from the preference evolution model of Alger et al. (2020) with the actor-centered representation of lineage fitness of Lehmann and Rousset (2020) to show that if genes delegate behavioral choice to the organism and information is incomplete, then the uninvadable behavior rule leads each organism to appear to be attempting to maximize a representation of personal inclusive fitness. Delegation of behavioral choice to the organism means that instead of being genetically determined, behavior is conceptualized as a choice of action or sequence of conditional actions—strategies—from a set of feasible strategies throughout lifespan. Behavioral choice is expected to evolve when the environment is complex or fluctuates within and between generations, making the genetic encoding of fixed behavioral responses inefficient (e.g., Stephens, 1991; Dugatkin, 2004; Dunlap and Stephens, 2009; Botero et al., 2015; Dridi and Lehmann, 2016). In fact, essentially all animals appear to learn actions (Dugatkin, 2004), enabling the organism to change behavior according to experience, thus as if achieving a choice among alternatives in a repertoire at some behavioral equilibrium. If, among alternatives, each individual chooses the strategy it prefers given the behavior of others, then it is as though individuals strive to maximize an objective or utility function. A profile of strategies forms a Nash equilibrium if no individual can increase its utility by unilaterally changing its strategy (e.g., Mas-Colell et al., 1995; Maschler et al., 2013).

The rest of this paper formalizes these concepts and is organized as follows. First, a model of pairwise assortative interactions is presented. Second, the uninvadability of specific behavior rules is derived. It is shown that a necessary condition for uninvadability is that a behavior rule entails Nash equilibrium play of a particular fitness game. Thereby, behavior rules can be restricted to utility maximization. Then, two utility functions that turn out to be uninvadable are contrasted: a semi-Kantian preference and a personal inclusive fitness preference.

## 2 Model

### 2.1 Biological assumptions

#### 2.1.1 Population structure and interactions

Consider a population of constant size with demographically homogeneous and asexual individuals (i.e., no effective age, stage, or sex structure). Individuals reproduce at discrete demographic time steps and pairwise assortative interactions occur among individuals within each generation. These interactions are assumed to arise due to limited genetic mixing, which may result from either (i) family structure, as in the original models of interaction between relatives (Hamilton, 1964; Michod, 1980), or (ii) spatial structure, as in the standard island model of limited dispersal (Wright, 1931; Rousset, 2004). The consequence of such assortation is that an individual will interact with other individuals from the same ancestral lineage; namely, identical-by-descent individuals, as well as individuals from the population at large, who belong to other lineages.

During pairwise interactions within a generation, each individual has access to the same set of strategies *χ* (Table 1 for a list of symbols), an element of which is defined as ‘a specification of what an individual will do in any situation in which it may find itself’ (Maynard Smith, 1982, p. 10). An interaction may thus involve repeated or multi-move activities, in which individuals can randomize over sequential actions (allowing for mixed strategies), and where the duration of the interaction may be uncertain. The strategies expressed by individuals in the population determine their survival and reproduction. Accordingly, 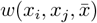 will denote the personal fitness; namely, the expected number of offspring (including the surviving self) produced over one demographic time step (or life cycle iteration) by individual *i* with strategy *x*_*i*_, when its interaction partner expresses strategy *x*_*j*_, in a population where the average strategy is 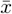.

**Table 1:** Definition of the symbols.

| Notations | Definitions |
| --- | --- |
| $\mathcal{X}$ | Set of strategies (assumed to be a compact and convex set in some topological vector space, see end of Appendix A for examples). |
| $\mathcal{U}$ | Set of continuous functions $u : \mathcal{X}^3 \rightarrow \mathbb{R}$. |
| $\Theta$ | Set of genetic traits or types. The nature of this set depends on what the traits code for. Under strategy evolution $\Theta = \mathcal{X}$, while under preference evolution $\Theta = \mathcal{U}$. |
| $b_\theta(x_s, x_d)$ | Behavior rule of type $\theta$ giving the set of acceptable strategies for its carrier given the strategy $x_s$ of lineage members and the strategy $x_d$ of non-lineage members. Formally, $b_\theta : \mathcal{X}^2 \rightrightarrows \mathcal{X}$ is a correspondence or set-valued function, whose range is a subset of $\mathcal{X}$ (Mas-Colell et al., 1995, p. 949). A behavior rule is an example of a choice rule (Mas-Colell et al., 1995, p. 10) applied to social interactions. |
| $\mathcal{X}_\theta$ | Set of strategy equilibria in a monomorphic $\theta$ population (eq. 1). |
| $\mathcal{B}_{\tau,\theta}$ | Set of strategy equilibria for a mutant-resident pair (eqs. 1–2). |
| $\mathcal{X}_{\text{NE}}$ | Set of Nash equilibria of the lineage fitness game when traits are strategies: $\Theta = \mathcal{X}$ (eq. 7). |
| $\arg \max_{x \in \mathcal{X}} f(x)$ | Best reply correspondence $\{x \in \mathcal{X} : f(y) \leq f(x) \text{ for all } y \in \mathcal{X}\}$ giving the set of maximizers of $f(x)$. |
| $W(\tau, \theta)$ | Invasion fitness (or lineage fitness or geometric growth ratio) of a mutant $\tau \in \Theta$ in a resident $\theta \in \Theta$ population under strategy equilibrium $(x^*(\tau, \theta), y^*(\theta)) \in \mathcal{B}_{\tau,\theta}$. |
| $\mathcal{W}(\tau, \theta)$ | Set of invasion fitness $W(\tau, \theta)$ stemming from all strategy equilibria $(x^*(\tau, \theta), y^*(\theta)) \in \mathcal{B}_{\tau,\theta}$. |
| $r(x^*(\tau, \theta), y^*(\theta))$ | Relatedness between a mutant and its partner under $(x^*(\tau, \theta), y^*(\theta)) \in \mathcal{B}_{\tau,\theta}$. |
| $u_{\theta_i}(x_i, x_j, \bar{x})$ | Personal utility of an individual $i$ with trait $\theta_i$ and strategy $x_i$ when its partner has strategy $x_j$ in a population with average strategy $\bar{x}$ ($\theta_i = \text{K}$ denotes the semi-Kantian preference, eq. 9, while $\theta_i = \text{H}$ is the inclusive fitness preference, eq. 10). Formally, $u_{\theta_i} : \mathcal{X}^3 \rightarrow \mathbb{R}$ and is assumed to be continuous (i.e., $u_{\theta_i} \in \mathcal{U}$). |
| $w(x_i, x_j, \bar{x})$ | Personal fitness: expected number of offspring (including the surviving self) produced over one demographic time step of an individual $i$ with strategy $x_i$ when its partner has strategy $x_j$ in a population with strategy $\bar{x}$. Formally, $w : \mathcal{X}^3 \rightarrow \mathbb{R}_+$, is assumed to be continuous, and constant population size entails that $w(x, x, x) = 1$ for all $x \in \mathcal{X}$. |
| $q(x_i, \bar{x})$ | Relative relatedness $r(x_i, \bar{x})/(1 + r(x_i, \bar{x}))$ of individual $i$ to its partner when $i$ has strategy $x_i$ and the average population strategy is $\bar{x}$. |
| $-c(x_i, x_j, \bar{x})$ | Average effect on own personal fitness of an actor with strategy $x_i$ when its partner has strategy $x_j$ in a population with average strategy $\bar{x}$ (eq. 11). |
| $b(x_i, x_j, \bar{x})$ | Average effect on the personal fitness of the interaction partner with strategy $x_j$ of an actor with strategy $x_i$ in a population with average strategy $\bar{x}$ (eq. 11). |

Each individual in the population is also endowed with a genetic trait (or type), which determines how it expresses its strategy. This trait is transmitted intact from parent to offspring, barring mutations. We will consider different kinds of trait spaces Θ, depending on the complexity of how individuals express their strategies (detailed in the next section). The overarching goal of the analysis is then to identify which traits from the set Θ are uninvadable when fixed in the population—that is, resistant to invasion by any mutant trait. Because of this focus on invasion analysis in a monomorphic population, there is no loss of generality in assuming that individual fitness depends on the average population strategy (rather than on the full distribution of strategies). In a monomorphic population, all individuals express the same genetic trait and are here assumed to express the same strategy.

#### 2.1.2 Behavior rules and information

In its most general form, an individual’s genetic trait determines a behavior rule—an internal mechanism that guides the choice of strategies from the set of alternatives *χ*. No assumptions are made regarding the complexity of the physiological or psychological processes underlying choice behavior, yet informational assumptions are imposed. This is because the game theory literature on the evolution of preferences (a special case of behavior rules, discussed below) demonstrates that the behavior rules favored by selection depend on the information individuals have about the behavior rules of others (e.g., Dekel et al., 2007; Heifetz et al., 2007a,b; Alger, 2023). Interactions are here assumed to occur under incomplete information: each individual’s behavior rule is its private information, meaning individuals are unaware of the genotype and behavior rule of others. This provides a useful baseline case for studying the evolution of behavior rules (and assessed in section 4).

More formally, the behavior rule *b*_*θ*_(*x*_s_, *x*_d_) for an individual with trait *θ* ∈ Θ is assumed to provide a subset of acceptable strategies to that individual from the set *χ*, given the strategy *x*_s_ of lineage members (i.e., individuals identical-by-descent to the focal *θ* carrier) and the strategy *x*_d_ of non-lineage members (i.e., individuals not identical-by-descent). To understand this concept, let us break it down (and see Table 1 for a formal definition). First, note that interactions often feature multiple possible behavioral outcomes. For instance, when individuals learn actions through reinforcement learning–by taking actions based on past rewards or penalties (Sutton and Barto, 1998)– multiple behavioral equilibria arise (e.g., Fudenberg and Levine, 1998; Dridi and Akçay, 2018). This is typical in coordination settings, which, essentially by definition, display multiple acceptable behavioral alternatives for an individual, but this feature holds more generally under social interactions (Binmore, 2007; Fudenberg and Tirole, 1991). Thus, a behavior rule, which is akin to the concept of a choice rule of decision theory (Mas-Colell et al., 1995, p. 10), should accommodate for multiple equilibria. This is captured by the behavior rule returning a set of acceptable strategies for its carrier. Second, under incomplete information, an individual does not know the strategy of its interaction partner, and may interact with both lineage and non-lineage members, owing to limited genetic mixing. Therefore, the behavior rule must also account for the strategy of each type of individual in relation to the focal individual. This is captured by the behavior rule depending on both the strategy of lineage and non-lineage members. The notion of a behavior rule is conceptually equivalent to that defined in Alger and Weibull (2012, p. 46), although they consider different informational assumptions.

### 2.2 Invasion analysis

In order to characterize which behavior rule is uninvadable, evolutionary invasion analysis is used (e.g., Eshel and Feldman, 1984; Metz et al., 1992; Ferrière and Gatto, 1995; Van Cleve, 2023) as applied to preference and behavior rule evolution (Alger et al., 2020; Alger and Lehmann, 2023). Consider then a population that is monomorphic for some resident behavior rule *b*_*θ*_ of type *θ* ∈ Θ, in which a mutant with behavior rule *b*_*τ*_ of type *τ* ∈ Θ arises and where Θ stands for the set of all feasible behavior rules. To characterize mutant uninvadability in this population, let us first focus on the monomorphic resident population and assume that all individuals use the same strategy *y*^*∗*^ at a behavioral equilibrium, to be called a resident strategy. This has to satisfy the fixed-point

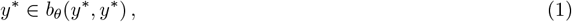

which says that the equilibrium resident strategy belongs to the set of acceptable strategies for a resident individual, given that all other individuals in the population, including lineage and non-lineage members, express the same strategy. Let *χ* _*θ*_ ⊂ *χ* denote the set of resident strategies satisfying eq. (1). When a mutant appears in such a population, all residents are assumed to express one and the same strategy. Given any such resident strategy *y*^*∗*^ ∈ *χ* _*θ*_, all mutants are assumed to use one and the same strategy, say *x*^*∗*^. This then has to satisfy the fixed-point

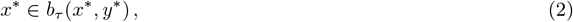

which says that the equilibrium mutant strategy belongs to the set of acceptable strategies for a mutant, given lineage members express the same strategy *x*^*∗*^ and non-lineage members express *y*^*∗*^. Denote by ℬ_*τ,θ*_ the (non-empty) set of equilibria satisfying the system (1)–(2) of fixed points. Any equilibrium pair (*x*^*∗*^, *y*^*∗*^) ∈ ℬ_*τ,θ*_ can be written as (*x*^*∗*^(*τ, θ*), *y*^*∗*^(*θ*)), where the dependence of strategies on the traits underlying the behavior rule is made explicit. This dependence enshrines the assumption of incomplete information: the resident strategy depends only on the resident trait (and thus behavior rule), while the mutant strategy depends on both mutant and resident traits.

It now follows that for any mutant *τ*, appearing in a single individual (the progenitor of the mutant lineage), in a resident population *θ* inducing behavioral equilibrium (*x*^*∗*^(*τ, θ*), *y*^*∗*^(*θ*)), the mutant goes extinct with probability one if and only if

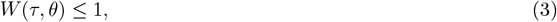

where

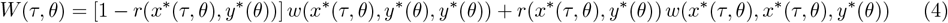

is the *lineage fitness* or *invasion fitness* of *geometric growth ratio* of the mutant (Alger and Lehmann, 2023, eqs. 1–2 under incomplete information). Lineage fitness can be read as the personal fitness of a randomly sampled mutant *τ* descending from the progenitor of the mutant lineage, averaged over the cases where the mutant interacts with a lineage member and those where it interacts with an individual from a different lineage (who is thus necessarily of type *θ*). The averaging depends on the pairwise relatedness *r*(*x*^*∗*^(*τ, θ*), *y*^*∗*^(*θ*)) between a mutant individual and its interaction partner. This is the probability that, conditional on an individual being of type *τ*, the interaction partner belongs to the same lineage and is thus of type *τ*. Namely, the probability that both individuals are identical-by-descent, meaning that their ancestral lineages coalesce in the progenitor of the lineage. Relatedness generally depends on the evolutionary process, and thus generally on the mutant and resident types and therefore on behavior (e.g., Mullon et al., 2016 and Box 1-2 of Alger and Lehmann, 2023). There are life-cycles, however, where relatedness is independent of mutant and resident types. For example, in a family-structured population, relatedness is determined solely by neutrality-based pedigree relationships, such as 1*/*2 for full siblings. This is implied by the model of Michod (1980), which also establishes that eq. (4) applies to sexual reproduction in family-structured populations in the absence of inbreeding, in which case *r* can simply be treated as a parameter.

Let us denote by 𝒲 (*τ, θ*) the set of lineage fitness resulting from all behavioral pairs (*x*^*∗*^(*τ, θ*), *y*^*∗*^(*θ*)) ∈ ℬ _*τ,θ*_. A behavior rule of type *θ* is said to be uninvadable if it is resistant to invasion by any behavior rule in the space of alternatives. That is, behavior rule *θ* is uninvadable if and only if

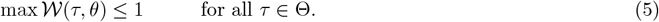

Uninvadability of the behavior rule *θ* thus preempts mutant invasion in the sense that the rule obtains weakly higher lineage fitness that any mutant can obtain under any behavioral equilibrium.

## 3 The genetical evolution of preferences

### 3.1 From behavior rules to preferences and Nash’s categorical imperative

In order to understand what type of behavior rules are favored by natural selection, it is useful to first examine behavior rules that induce commitment to one particular strategy. In that case, the strategy of an individual can be conceived to be directly genetically determined, a case henceforth called strategy evolution (consistent with previous terminology, e.g., Alger, 2023). Under strategy evolution, all individuals with behavior rules in the set Θ each have a unique strategy. We can thus identify traits by their strategy and set Θ = *χ*. Then, for a mutant strategy *τ* = *x* and resident strategy *θ* = *y*, it follows from eqs. (3)–(5) that max 𝒲 (*x, y*) = *W* (*x, y*), where *W* (*x, y*) = [1 − *r*(*x, y*)] *w*(*x, y, y*)+*r*(*x, y*) *w*(*x, x, y*) is the lineage fitness of trait *x* in population *y*. Thus, a necessary and sufficient condition for strategy *y*^*∗*^ ∈ *χ* to be uninvadable is that it preempts mutant invasion by obtaining the maximal lineage fitness that can be obtained in a population where all individuals express *y*^*∗*^:

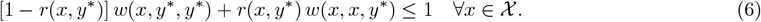

Equivalently, strategy *y*^*∗*^ is uninvadable if and only if it is a best reply to itself in terms of lineage fitness, meaning that

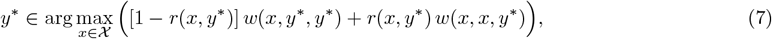

where the right-hand side means the set of traits maximizing lineage fitness (see also Table 1). In other words, *y*^*∗*^ is a symmetric Nash equilibrium, where lineage fitness cannot be increased by unilaterally changing mutant trait value. Denote by *χ* _NE_ the set of such equilibria (assumed non-empty, see Appendix A for conditions thereof). In terms of these concepts, Appendix B shows the first result of this paper.

#### Result 1.

*A behavior rule b*_*θ*_ *of type θ* ∈ Θ *is uninvadable if and only if χ* _*θ*_ ⊆ *χ* _NE_.

This generalizes a result by Alger et al. (2020, Proposition 1) and states that for a behavior rule to be uninvadable, it must induce resident equilibria that are uninvadable strategies, and thus constitute Nash equilibria of the lineage fitness game. Nash equilibrium play by individuals is therefore not an assumption of the model, but a categorical imperative–a binding principle–imposed by evolution for a behavior rule to be uninvadable. This allows one to restrict attention to behavior rules that entail utility maximization since this induces Nash equilibrium play (and Appendix A recalls that an individual’s utility also provides a representation of its preference).

Let us thus denote by 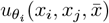 the personal utility of individual *i* with trait *θ*_*i*_ expressing strategy *x*_*i*_ when its interaction partner expresses strategy *x*_*j*_ in a population where the average strategy is 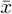. Under incomplete information, an individual does not know the behavior rule and thus strategy of its interaction partner, since this information is private. The information assumed here to be available to each individual is the relatedness to its interaction partner, since this is the default information that should be accumulated by the evolutionary process (Frank, 1998, pp. 95 and 104). In this case, the realized utility of an individual is given by an expectation of its utility over the case where the interaction partner is a lineage member, which occurs with a probability given by relatedness, and a non-lineage member, with complementary probability.

Under incomplete information, each individual thus maximizes its expected utility and the resulting Nash equilibrium is said to be Bayesian (Mas-Colell et al., 1995; Fudenberg and Tirole, 1991). When a mutant with utility *u*_*τ*_ interacts with its partner in a population where residents have utility *u*_*θ*_, the strategy pair (*x*^*∗*^(*τ, θ*), *y*^*∗*^(*θ*)) is a (Bayesian) Nash equilibrium if (a) *y*^*∗*^ is a preferred strategy for the resident, given that other residents use *y*^*∗*^; and (b) *x*^*∗*^ is a preferred strategy for a mutant, given that other mutants use *x*^*∗*^ and residents use *y*^*∗*^, and given that the mutant applies the probability *r*(*x*^*∗*^, *y*^*∗*^) of being matched with another mutant, when all individuals play their equilibrium strategies (Alger et al., 2020). Thus, the Nash equilibrium (*x*^*∗*^, *y*^*∗*^) between the mutant and the resident is characterized by the system (1)–(2) of fixed points with resident and mutants behavior rules given, respectively, by

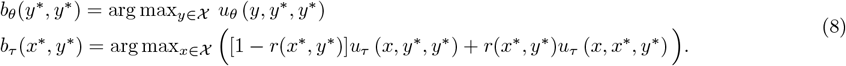

Each behavior rule thus provides a set of best replies in terms of expected utility, given the equilibrium strategies of lineage and non-lineage members. Note that the set *χ*_*θ*_ of resident equilibria under *u*_*θ*_ are thus all Nash equilibria satisfying *y*^*∗*^ ∈ arg max_*y∈ χ*_ *u*_*θ*_ (*y, y*^*∗*^, *y*^*∗*^) and the utility therein can be expanded as *u*_*θ*_ (*y, y*^*∗*^, *y*^*∗*^) = [1 − *r*(*y*^*∗*^, *y*^*∗*^)]*u*_*θ*_ (*y, y*^*∗*^, *y*^*∗*^) + *r*(*y*^*∗*^, *y*^*∗*^)*u*_*θ*_ (*y, y*^*∗*^, *y*^*∗*^), making explicit that a resident individual interacts with lineage and non-lineage members as well.

By now restricting the set of evolving traits Θ to the set of continuous utility functions, we can ask what is the form of the uninvadable personal utility?

### 3.2 Preference evolution and Kant’s categorical imperative

Consider first the utility of type K ∈ Θ defined by

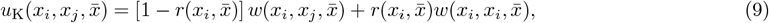

which is to be interpreted as follows. The first fitness term is the focal individual’s realized personal fitness, given the strategies used in the population. The second fitness term is the personal fitness that the individual would realize if – hypothetically – the interaction partner used the same strategy *x*_*i*_, instead of its own strategy *x*_*j*_. The individual thereby evaluates what would happen if the interaction partner were to follow the same course of action as self. This second fitness term captures a form of the first formulation of Kant’s categorical imperative: “act as if the maxims of your action were to become through your will a universal law of nature” (Kant, 1785). The relatedness 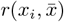that appears in the weighing of each term depends on the focal’s trait *x*_*i*_ and the average 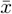 in the population. With this, we are led to the next result of this paper shown in Appendix C.

#### Result 2.

*The utility function u*_K_ *is uninvadable*.

In an uninvadable population state of behavior rule evolution, it is thus as if each individual appears to behave to attempt to maximize the semi-Kantian preference, given that each other individual in the population does the same. Intuitively, under incomplete information, it is as if the evolutionary process favors modes of internal information processing that are analogs of the evolutionary process itself, since the function of strategies is to bring forth fitness maximization, and the information accessed by behavior rules is analogous to that as if an evolutionary process was to occur at the level of strategies.

The uninvadabilty of *u*_K_ generalizes a result by Alger et al. (2020, their Proposition 2, see Appendix C for comparison). In the absence of limited genetic mixing and social interactions where lineage fitness can be written in the multiplicatively separable way *W* (*τ, θ*) = *F* (*τ*) *G*(*θ*) for some functions *F* and *G* depending, respectively, only on mutant and resident traits, the uninvadabilty of *u*_K_ is implied by the results of Cooper (1987, 2001). He further showed that a behavior rule that is uninvadable under these conditions is Savage rational and thus entails maximization of expected utility over all within generation uncertainty. Since personal fitness *w* resolves all within generation uncertainty, *u*_K_ can, under the present model, *defacto* be interpreted as an expected utility over within generation uncertainty.

### 3.3 Actor-centered preference and Hamilton’s categorical imperative

Consider now the utility of type H ∈ Θ defined by

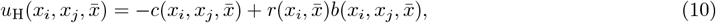

where 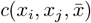 is the personal cost to the actor with strategy *x*_*i*_ when the interaction partner has strategy *x*_*j*_ in a population with average strategy 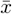, while 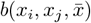 is the benefit to the interaction partner, which is the recipient of the actor’s strategy. These costs and benefits satisfy

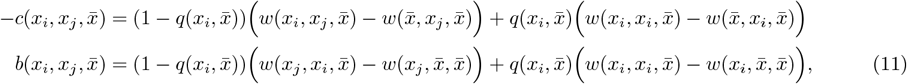

where

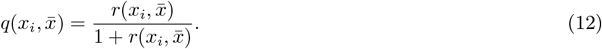

In the first line of eq. (11), the interpretation of 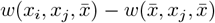 is the actor’s fitness gain of expressing its strategy *x*_*i*_ instead of 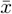, given the partner’s strategy is *x*_*j*_. Likewise, 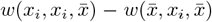 is the actor’s fitness gain of expressing strategy *x*_*i*_ instead of 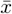, given the partner’s strategy is *x*_*i*_. Hence, 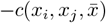 can be thought of as the average effect on the fitness of the actor expressing its strategy *x*_*i*_ instead of hypothetically acting as an average individual in the population, and on the supposition that the interaction partner will play the same strategy as self, with probability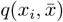, which is relative relatedness. In the second line, 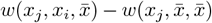 is the fitness gain to a recipient with strategy *x*_*j*_ resulting from the actor expressing strategy *x*_*i*_ instead of 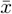, while 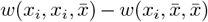 is the fitness to a recipient expressing strategy *x*_*i*_ resulting from the actor expressing strategy *x*_*i*_ instead of 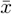. Hence, 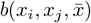 measures the average effect on the fitness of the interaction partner of the actor expressing its own strategy instead of hypothetically acting as an average individual in the population, and on the supposition that the interaction partner will play the same strategy as self with probability 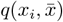.

The utility function *u*_H_ can be interpreted as an actor-centered preference, since it associates the consequence of own behavior with the affected individuals (Rousset, 2004, Fig.7.1), that is self, the interaction partner, and others in the population (who have zero relatedness to the focal actor). This is the essence of the inclusive fitness approach (Hamilton, 1970; Rousset, 2004), and so *u*_H_ can be regarded as a personal inclusive fitness (strictly speaking a personal inclusive fitness effect, since a “1” should be added to yield inclusive fitness 1 − *c* + *b*, but this does not alter any conclusion). If the average effects 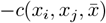 and 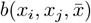 were the results of a gene substitution (instead of a behavioral substitution between *x*_*i*_ and 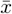), then the interpretation of *average effects* would literally be that of Fisher (1930, 1941) obtained from the least squares method. In the behavioral context, the personal inclusive fitness eq. (10) can thus be regarded as the best additive system for associating own behavior with the individuals affected from the actual behaviors expressed by interacting individuals. It is this additivization that gives rise to the special form of the weight in eq. (12) (see Lehmann and Rousset, 2020, Appendix C.3 for a derivation of eq. (11) using least-squares arguments). The additivization also generates the hallmark feature of eq. (10), namely Hamilton’s binding principle: each individual should treat recipients (including self) according to their degree of genetic relatedness to self. This, in turn, implies the necessity of positive relatedness for the expression of personal self-sacrifice that benefits others. Indeed, when *r* = 0, *u*_H_ = −*c*, where *c* is the average effect of the actor’s behavior on its own fitness, and *c >* 0 for any behavior *x*_*i*_ that decreases personal fitness relative to the expression of some self-preserving behavior 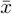.

The actor-centered preference *u*_H_ satisfies the same resident equilibria as the semi-Kantian preference *u*_K_, since utility maximization is unaffected by affine transformations (e.g., Mas-Colell et al., 1995; Maschler et al., 2013) and (see Appendix D). In force of the uninvadability of *u*_K_, we thus immediately obtain the last result of this paper.

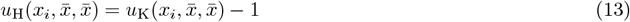

#### Result 3

*The utility function u*_H_ *is uninvadable*.

Hence, in an uninvadable population state of behavior rule evolution, it is as if each individual appears to attempt to maximize its personal inclusive fitness, given that each other individual in the population does the same.

## 4 Discussion

This paper demonstrates the existence of a model that satisfies the maxim of the introductory quote for interactive social behavior of arbitrary complexity. The maximization of personal inclusive fitness occurs in the sense that, when confronted with a choice among alternatives, an individual strives to increase the sum of a self-regarding interest and a relatedness-weighted other-regarding interest. Both interests involve evaluating the consequences of one’s own behavior, rather than expressing average population behavior, on a statistical average fitness to the recipient (eq. (11)). This complexity aligns with the general formulation of inclusive fitness in population genetics, which necessarily entails a comparison of fitness effects, even in the absence of behavioral choice (Gardner et al., 2011; Rousset, 2015; Lehmann and Rousset, 2020). An alternative—and for some, perhaps more intuitive—representation of an individual’s preferences is given by the semi-Kantian preference, eq. (9), which is also uninvadable. This representation is popular in the game theory literature (Bergstrom, 1995; Binmore, 1998; Alger and Weibull, 2013). Both the inclusive fitness and semi-Kantian preference representations are as general as the lineage fitness of the model, and thus, no particular restrictions have been placed on them.

Previous work attempting to identify conditions under which personal inclusive fitness is maximized in an evolutionary setting has focused on the case of strategy evolution, where maximization can be equated with uninvadability. The bulk of this work, however, is restricted to cases where fitness depends additively on the traits of the focal individual and its interaction partners (e.g., Grafen, 2006; Okasha et al., 2014; Lehmann and Rousset, 2014), such that best responses to population traits, and thus Nash equilibria, play no role. This rules out most forms of social interactions. In work acknowledging behavioral complexity and Nash equilibrium play, skepticism has been expressed regarding the possibility of obtaining a genuine personal representation of inclusive fitness (Lehmann et al., 2015; Lehmann and Rousset, 2020). The difficulty is that relatedness in lineage fitness depends on individuals’ strategies (eq. 4), which implies that relatedness will also depend on one’s own strategy in personal inclusive fitness (eq. 10). Yet, the actual genetic relatedness between actor and recipient depends on the population genetic process, and thus on the behavior of ancestral, identical-by-descent lineage members, not directly on the actor’s own behavior. On the other hand, if preferences are meant to represent individual behavior, whether relatedness is a real property of an individual or of its lineage can be put aside, since what matters is that “inclusive fitness is that property of an individual organism which will appear to be maximized when what is really being maximized is gene survival” (Dawkins, 1978, p. 73). From this perspective, the uninvadable personal inclusive fitness preference is genuine.

The positive results of this paper were derived under the assumption of incomplete information, where the available information about others is represented by the average relatedness between interacting individuals. While this assumption is strong, it corresponds to the default working assumption in kin selection reasoning (Frank, 1998, Chapter 6). As described by Frank (1998, Chapter 6), individuals may acquire more fine-grained information through various mechanisms of discrimination. In the present context, they may thus obtain information about each other’s behavioral rules—potentially even to the extent of knowing each other’s preferences—in which case the information is said to be complete (see Akçay and Van Cleve, 2012; Alger and Weibull, 2012; Alger and Lehmann, 2023 for models of preference evolution under assortative interactions with complete information). While it is more realistic that individuals possess information that lies somewhere between these two special cases (see Heifetz et al., 2007a,b for a formalization under panmixia), the appeal of the baseline case of incomplete information lies in two key aspects. It enables the derivation of preference representations that (i) are interpretable, aligning with biological fitness, and (ii) are transferable across different social interaction settings (or games), unlike the other informational settings where predictions become interaction specific. This, in turn, allows for the formulation of general behavioral predictions (which appear consistent with empirical findings, Levy and Lo, 2022; van Leeuwen and Alger, 2024). Further modeling work could explore the genetic evolution of preferences under various informational settings, examine adaptive polymorphism in preferences, consider class-structured populations, and incorporate complexity costs into decision-making behavior.

In conclusion, starting with the demographic concept of resistance to invasion, the analysis shows that, at an evolutionary equilibrium of behavior rule evolution under incomplete information, individuals must behave as if their lifetime cumulative behavior were goal-directed. This goal can be interpreted as nepotistic, involving two differences in expected fitness: a cost to the actor and a benefit to the recipient, weighted by their relatedness to the actor. This result holds regardless of the strength of selection on behavioral rules or the complexity of behavioral interactions, with moment-to-moment actions and reactions potentially being only secondary, tertiary, or even more distant correlates of survival and reproduction. The model thus provides formal support for long-standing principles in behavioral ecology (Davies et al., 2012, p. 312), as well as for the perspective on human behavior developed by Alexander (1979), which merits renewed attention and careful consideration.

## Acknowledgments

I thank the associate editor and an anonymous reviewer for their valuable comments, which greatly helped to clarify the presentation of the paper. I’m especially grateful to another reviewer for pointing out a quirk in a previous version of a proof, the correction of which led to a fundamental strengthening of the results. Finally, I thank Ingela Alger and Jörgen Weibull for discussions.

## Appendix A Utility functions, preferences, and Nash equilibrium

We here first recall the standard relationship between the concept of a utility function and the concept of a preference, and then recall sufficient conditions for existence of a Nash equilibrium.

For any pair of tuples of strategies, 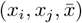 and 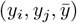, the preference ordering ⪰_*i*_ of individual *i* over own, interaction partner *j*, and population average strategies determines whether the individual prefers 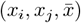, or is indifferent between both. When such a preference ordering is complete and transitive, it can be described by a function that assigns a real number to each strategy combination, the utility function *u*_*i*_ of individual *i*. Namely, there exists a continuous function *u*_*i*_ : *χ* ^3^ → ℝ satisfying 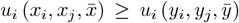 if and only if 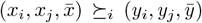 (see, e.g., Osborne and Rubinstein, 1994; Mas-Colell et al., 1995; Binmore, 2011; Maschler et al., 2013 and Appendix A). A symmetric Nash equilibrium *x*^*∗*^ is a profile of strategies with the property that (*x*^*∗*^, *x*^*∗*^, *x*^*∗*^) ⪰_*i*_ (*x*_*i*_, *x*^*∗*^, *x*^*∗*^) for all *x*_*i*_ ∈ *χ* and all individuals in the population. This can be restated as *x*^*∗*^ ∈ arg max_*x∈X*_ *u*_*i*_(*x, x*^*∗*^, *x*^*∗*^).

Letting U be the set of continuous functions *u* : *χ* ^3^ → ℝ, each individual in the population then has a preference that admit representation by some function *u* ∈ 𝒰. In the evolutionary setting of the main text under incomplete information, each individual chooses its strategy so as to maximize the expected value of its utility function, under the assumption that the interaction partner’s strategy choice is similar to self with probability given by relatedness (see eq. 8). The utility function itself is determined by the genetic trait of the individual, and we write *u*_*θ*_ for the utility function of type *θ*. Under preference evolution the set of traits, or types Θ, can then be taken as the set of utility functions: Θ = 𝒰.

Le us now turn to describe the standard sufficient condition for the existence of a Nash equilibrium in a game with continuous payoff function *π* : *χ* ^*n*^ → R, where *n* is the numbrer of “players” (e.g., Osborne and Rubinstein, 1994; Mas-Colell et al., 1995; Fudenberg and Tirole, 1991; Carmona, 2013). When *n* = 2, *π*(*x, y*) can be taken to be the invasion fitness *W* (*x, y*) in section 3.1, while when *n* = 3, *π*(*x, y, z*) can be taken to be the utility function *u*(*x, y, z*). A symmetric Nash equilibrium *x*^*∗*^ satisfies the fixed point

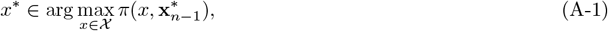

where 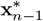 is the (*n* − 1) dimensional vector with all components *x*^*∗*^. Such a Nash equilibrium exists (i.e., the set of best-responses satisfying eq. (A-1) is non empty) if the payoff function *π* is quasi-concave in its first argument and the strategy set *χ* is a compact and convex subset of a topological vector space (Carmona, 2013, Theorem 2.1, p. 6). Examples of topological vector space include (i) the Euclidean space (i.e., ℝ ^*d*^ with *d* a positive integer), which allows to model continuous action spaces or mixed strategies over a finite set of actions; (ii) the space of sequences of real numbers (i.e., ℝ ^ℕ^), which allows to model streams of actions under repeated games where the duration of the interaction may be uncertain; (iii) normed vector spaces like the space of continuous functions on a closed interval, which allows to model life-history strategies.

## Appendix B Proof of result 1

We here show that a behavior rule *b*_*θ*_ of type *θ* ∈ Θ is uninvadable if and only if *χ* _*θ*_ ⊆ *χ* _NE_, i.e., the set of resident equilibria satisfying eq. (1) belongs to the set of equilibria satisfying eq. (7) in an uninvadable population state. For this, let us first observe that under behavior rule evolution it follows by combining eq. (4) and eq. (5) that the condition for a behavior rule of type *θ* ∈ Θ to be uninvadable can be expressed in the form:

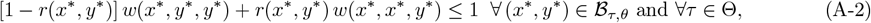

where the shorthand *x*^*∗*^ = *x*^*∗*^(*τ, θ*) and *y*^*∗*^ = *y*^*∗*^(*θ*) is used throughout this section.

Let us now show that *χ* _*θ*_ ⊆ *χ* _NE_ is a sufficient condition for behavior rule of type *θ* to be uninvadable. Suppose that *χ* _*θ*_ ⊆ *χ* _NE_. Then for each *y*^*∗*^ ∈ *χ*_*θ*_, [1 − *r*(*x, y*^*∗*^)] *w*(*x, y*^*∗*^, *y*^*∗*^) + *r*(*x, y*^*∗*^) *w*(*x, x, y*^*∗*^) ≤ 1 ∀*x* ∈ *χ*; namely, eq. (6) is satisfied for any strategy *x* ∈ *χ* played by mutants. In other words, there exists no *τ* ∈ Θ for which some (*x*^*∗*^, *y*^*∗*^) ∈ ℬ_*τ,θ*_ does not satisfy the inequality in eq. (A-2). Hence, the condition in eq. (A-2) for *θ* to be uninvadable in Θ is satisfied.

Let us next show that *χ*_*θ*_ ⊆ *χ*_NE_ is a necessary condition for a behavior rule of type *θ* to be uninvadable. Suppose to the contrary that *θ* is uninvadable and there exists some resident equilibrium *y*^*∗*^ ∈ *χ*_*θ*_ such that *y*^*∗*^ ∉ *χ*_NE_. Then, there exists some strategy *x* ∈ *χ* for which [1 − *r*(*x, y*^*∗*^)] *w*(*x, y*^*∗*^, *y*^*∗*^) + *r*(*x, y*^*∗*^) *w*(*x, x, y*^*∗*^) *>* 1 (i.e., eq. 6 is not satisfied when *y*^*∗*^ ∉ *χ*_NE_). Consider a mutant behavior rule of type *τ* that induces mutants to play this same strategy, namely *x*^*∗*^ = *x*, whatever the strategy expressed by the resident. Then, there exists (*x*^*∗*^, *y*^*∗*^) ∈ ℬ_*τ,θ*_ for which eq. (A-2) is not satisfied. This means that the behavior rule of type *θ* is invadable, which contradicts the supposition.

## Appendix C Proof of result 2

We here show that the utility function *u*_K_ is uninvadable. Consider any Nash equilibrium strategy *y*^*∗*^ ∈ *χ*_NE_ such that eq. (7) holds. To see that this strategy *y*^*∗*^ is also a resident strategy under the semi-Kantian preference *u*_K_, note that from eq. (9), *y*^*∗*^ is a resident strategy under *u*_K_ if and only if

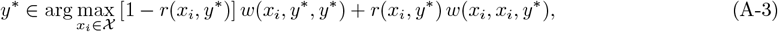

which indeed satisfies *y*^*∗*^ ∈ *χ* _NE_. To see this, compare eq. (A-3) to eq. (7), which are the same, up to the labeling of the dummy variable which is inconsequential. This together with Proposition 1 implies that *u*_K_ is uninvadable.

This uninvadability result of *u*_K_ is similar but stronger than Proposition 2 of Alger et al. (2020), who show that *u*_K_ is uninvadable if the set of resident equilibria *χ*_K_ under *u*_K_ is singleton. The reason for this difference is that Alger et al. (2020) defined the evolving utility function of interacting individuals to depend only on the strategy of patch members, and not the population strategy; that is 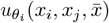 is reduced to *u*_*θ*_*i* (*x*_*i*_, *x*_*j*_) for player *i* interacting with player *j*. Hence, there is no concept of background population. This then allows for resident equilibria under *u*_K_ to differ from that of an uninvadable strategy (Alger et al., 2020, eq. 55).

## Appendix D Proof of eq. (13)

Substituting eq. (12) into eq. (11), setting 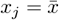, and using the property 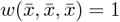 (see Table 1) yields

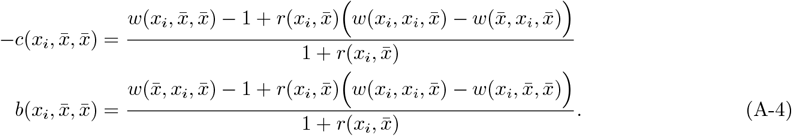

Thus

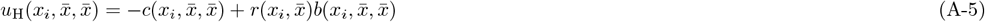

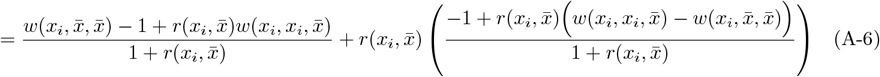

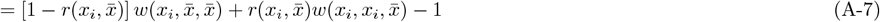

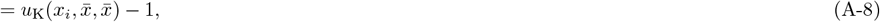

where the second equality is obtained by eliminating 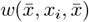.

